# Relative evolutionary rates in proteins are largely insensitive to the substitution model

**DOI:** 10.1101/304758

**Authors:** Stephanie J. Spielman, Sergei L. Kosakovsky Pond

**Affiliations:** Institute for Genomics and Evolutionary Medicine, Temple University, Philadelphia, PA 19122.

**Keywords:** evolutionary rates, model selection, protein evolution, phylogenetics

## Abstract

The relative evolutionary rates at individual sites in proteins are informative measures of conservation or adaptation. Often used as evolutionarily-aware conservation scores, relative rates reveal key functional or strongly-selected residues. Estimating rates in a phylogenetic context requires specifying a protein substitution model, which is typically a phenomenological model trained on a large empirical dataset. A strong emphasis has traditionally been placed on selecting the “best-fit” model, with the implicit understanding that suboptimal or otherwise ill-fitting models can potentially bias inferences. However, the pervasiveness and degree of such bias has not been systematically examined. We investigated how model choice impacts site-wise relative rates from a large set of empirical protein alignments. We compared models designed for use on any general protein, models designed for specific domains of life, and the simple equal-rates Jukes Cantor-style model (JC). As expected, information theoretic measures showed overwhelming evidence that some models fit the data decidedly better than others. By contrast, estimates of site-specific evolutionary rates were impressively insensitive to the substitution model used, revealing an unexpected degree of robustness to potential model misspecification. A deeper examination of the fewer than 5% of sites for which model inferences differed in a meaningful way showed that the JC model can uniquely identify rapidly-evolving sites that models with empirically-derived exchangeabilities fail to detect. We conclude that relative protein rates appear robust to the applied substitution model, and any sensible model of protein evolution, regardless of its fit to the data, should produce broadly consistent evolutionary rates.

## Introduction

That the rates of substitution are not constant across a sequence, and are modulated by a multitude of processes and forces has been recognized since the dawn of modern comparative evolutionary analysis (Uzzell and Corbin 1971; Echave et al. 2016). Failing to account for site-to-site rate heterogeneity would be considered a neophyte error in contemporary applications, since it can lead to biased parameter estimation or incorrect phylogenies (e.g., Yang 1996). In addition to being an important confounder that needs to be corrected for, the distribution of rates across sites is of essential interest in its own right in many applications. As an example, the majority of analyses that seek imprints of natural selection on sequence data do so by inferring and interpreting the distributions of synonymous and non-synonymous substitution rates and interpreting their properties (e.g., Delport et al. 2009).

In the context of protein sequence analysis, low site-specific substitution rates have served as a proxy for evolutionary conservation, and similarly high rates have been regarded as a correlate of adaptation or positive selection (Sydykova and Wilke 2017). Positions which play key roles in protein functions, including those involved in protein-protein or protein-ligand interactions or those at or near active regions, tend to evolve very slowly and are highly conserved (Jack et al. 2016; Echave et al. 2016; Sydykova et al. 2018). By contrast, sites will tend to evolve rapidly if they interact with rapidly changing external stimuli, for example if they are involved in chemosensory activity (Spielman and Wilke 2013; Almeida et al. 2015), or mediate immunity, i.e. pathogen surface proteins or key regions of host immune genes (Tusche et al. 2012). Indeed, searching for conserved immune epitopes is a popular approach to finding vaccine or drug targets (Garcia-Boronat et al. 2008).

Evolutionary rates at individual sites in proteins are commonly measured as *relative* quantities, that indicate how quickly the site evolves relative to the “mean” protein rate. In such approaches, empirically-derived models of protein evolution are fit to data in a phylogenetic framework, using either maximum-likelihood or Bayesian methods (Pupko et al. 2002; Mayrose et al. 2004; Fernandes and Atchley 2008; Nguyen et al. 2015; Ashkenazy et al. 2016; Spielman and Kosakovsky Pond 2018). Available substitution models can be loosely dichotomized into two classes. “Generalist” models are inferred from training data that comprise proteins from many domains of life, e.g., JTT (Jones et al. 1992) or LG (Le and Gascuel 2008), and are meant to represent the evolutionary predilections of many different proteins. “Specialist” models, by contrast, are trained on data aimed to capture the properties of a particular taxonomic or biological group, e.g., mtREV for mammalian mitochondrial sequences (Adachi and Hasegawa 1996) or AB for human antibody sequences (Mirsky et al. 2015).

Because site-specific rates are usually estimated by conditioning on a specific model of sequence evolution, one can reasonably assume that the estimated distribution of rates among sites will be influenced by the choice of the evolutionary model, which can be considered a nuisance parameter when rate estimates are of primary interest. It comes as no surprise that the question of model selection has assumed a prominent role in evolutionary rate inference, with popular implementations, such as *ProtTest* (Darriba et al. 2011) garnering thousands of citations.

Historically, protein substitution and similarity scoring models have primarily been developed and benchmarked in the specific context of homology identification (Henikoff and Henikoff 1992) and phylogenetic reconstruction (Le and Gascuel 2010). As a consequence, the relative performance of these models for inferring evolutionary rates from protein sequences has not been extensively studied. When this facet of model performance is mentioned, it is usually done in passing. For example, Landau et al. (2005) suggested that rates inferred with different models may be similar but feature “non-negligible” differences.

We undertook a systematic comparison of site-specific rate distributions inferred using three generalist models, three specialist models, and the Jukes-Cantor equal-rates model (also known as JC, Jukes and Cantor 1969). As expected, standard likelihood-score based information criteria used in phylogenetic model selection revealed very strong model preferences for all alignments. Contrary to prevailing expectation, we found that across alignments ranging in taxonomic scope and levels of sequence divergence, models yielded rates that were nearly perfectly correlated. Even when we deliberately misapplied a specialist model, e.g., by using a mitochondrial model on chloroplast data, we obtained rates that were almost perfectly correlated with the rates inferred under the cognizant specialist model. It was only when using the extreme case of the equal-rates JC model that yielded, albeit rarely, rates meaningfully different from models with unequal residue exchangeabilities.

Our results imply that some features of the evolutionary inference are quite robust to model misspecification, and further any sensible model of protein evolution is likely to produce largely consistent evolutionary rate patterns in many settings. One one hand, this finding is not as surprising as it may appear, because many evolutionary-rate analyses analyses have been reported robust to various severe modeling violations at least based on simulated data and practical use cases [e.g., Anisimova et al. (2001)]. On the other hand, our results suggest that, depending on what estimates are of primary interest, standard model selection approaches may be suboptimal, since they do not identify the source of improvement in fit. In particular, our finding suggests that for some important applications, it is not necessary to waste CPU cycles on exhaustive model selection, and that alternative evaluative measures of goodness-of-fit could be more informative about the impact of evolutionary model choice on interpretable parameter inference (Brown 2014).

## Methods

### Data Collection and Processing

We collected alignments from four distinct classes of proteins: enzymes, Metazoan mitochondrial data, green land plant chloroplast data, and mammalian G protein-coupled receptor data (GPCRs).

For the enzyme dataset, we randomly selected 100 alignments with at least 25 unique sequences and corresponding phylogenies from Jack et al. (2016) for analysis. We prepared the mitochondrial and chloroplast datasets each as follows. First, we compiled a list of complete Metazoan mitochondrial and green land plant genomes from the NCBI genomes database, of which there were 7515 and 1026, respectively. For each dataset, we randomly chose 350 taxa to include in analysis. For these taxa, we obtained all genomic protein sequences from NCBI. After discarding sequences with amino acid ambiguities, and retaining only genes for which at least 100 taxa contained full-length sequences, we made gene-specific alignments using `MAFFT` v7.305b (Katoh and Standley 2013) and inferred phylogenies using `FastTree2` (Price et al. 2010) with the LG+F substitution model (Le and Gascuel 2008).

Finally, we retrieved 227 alignments of mammalian GPCRs analyzed by Spielman and Wilke (2013), filtered to include at least 20 sequences. We reconstructed phylogenies for these alignments using FastTree2 (Price et al. 2010) with the LG+F model (Le and Gascuel 2008), as the original phylogenies had been constructed using masked alignments with reduced information.

### Rate inference

We then inferred relative protein evolutionary rates with the LEISR (Spielman and Kosakovsky Pond 2018; Pupko et al. 2002) method in HyPhy version 2.3.8 (Kosakovsky Pond et al. 2005), using seven different amino-acid evolutionary models. We used three generalist models: LG (Le and Gascuel 2008), WAG (Whelan and Goldman 2001), JTT (Jones et al. 1992), three specialist models: mtMet (Metazoan mitochondrial, Le et al. 2017), gcpREV (green-plant chloroplast, Cox and Foster 2013), and HIVb (between-host HIV-1, Nickle et al. 2007), and the equal-rates JC model (Jukes and Cantor 1969). We inferred rates under each model with (+*G*) and without a four-category discrete gamma distribution to model site heterogeneity during branch length optimization. All inferences used empirical (+*F*) equilibrium residue frequencies. We processed LEISR output for subsequent analysis using the Python helper package phyphy (Spielman 2018). All analyses interpreted the raw relative rates returned by LEISR, i.e. rates were neither normalized nor standardized in any way. In addition, we right-censored any inferred rate with *MLE* ≥ 1000 to all have the value *MLE* = 1000, because this value is effectively the numerical infinity in this context. We quantified estimation error of individual retaliative rates using profile likelihood, tabulating approximate 95% confidence intervals, using the critical values of the 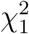 distribution.

### Statistical Analysis Availability

Data analysis was primarily conducted in the R programming language (R Core Team 2017), with substantial use of the `tidyverse` (Wickham 2017) suite of data analyis and visualization tools. All code and data associated with this work is freely available from the GitHub repository https://github.com/sjspielman/protein_rates_models.

## Results

We compiled 419 alignments selected to represent both “general” and “specialist” proteins: a dataset of enzyme alignments randomly selected from Jack et al. (2016), a dataset of mammalian G protein-coupled receptor (GPCR) alignments from Spielman and Wilke (2013), a dataset of green land plant chloroplast alignments, a dataset and Metazoan mitochondrial alignments (see Methods for details). Table 1 provides an overview of each dataset’s characteristics.

**Table 1:** Dataset used for rate estimation.

| Dataset | Class | $N^*$ | Median sites (IQR) | Median sequences (IQR) | Median tree length<sup>†</sup> (IQR) |
| --- | --- | --- | --- | --- | --- |
| Enzyme | Generalist | 100 | 564 (371) | 301 (104) | 82.4 (34.0) |
| Mitochondria | Specialist | 13 | 464 (357) | 344 (2) | 86.2 (49.4) |
| Chloroplast | Specialist | 79 | 238 (328) | 341 (15) | 7.9 (7.52) |
| GPCR | Generalist | 227 | 385 (156) | 22 (4) | 1.6 (1.75) |

\*: Number of alignments in the dataset.

^*^: Number of alignments in the dataset.

^†^: Tree length is computed as the sum of branch lengths, measured in expected substitutions per site, from the phylogeny (all built under the LG+F model) used as LEISR input. See Figures S1-2 for details on how models affect tree lengths.

We inferred site-specific relative evolutionary rates from each alignment under three generalist models (LG, WAG, JTT), three specialist models (mtMet, gcpREV, HIVb), and the JC model with equal rates. Respectively, these three specialist models were originally trained on Metazoan mitochondrial sequences (Le et al. 2017), green plant chloroplast sequences (Cox and Foster 2013), between-host HIV-1 (subtype M) sequences (Nickle et al. 2007).

We inferred rates with each model twice, with and without including a gamma distributed component to model rate heterogeneity during the branch length optimization step, producing fourteen sets of relative rate estimates per alignment. Throughout, we use +*G* to refer to a model which incorporated gamma-distributed rate heterogeneity. For a given alignment, rates inferred with any model employed the same equilibrium frequencies, empirically derived from the alignment, but a different exchangeability matrix. Any differences we identified between model inferences, therefore, were directly attributable to the differences in amino-acid exchangeabilities between model matrices. As such, we use the terms “model” and “matrix” interchangeably. We use the term “MLE” to refer to maximum-likelihood point estimates of relative rates.

*A priori*, we expected that gcpREV should best fit alignments from the chloroplast dataset, and that mtMet should best fit alignments from the mitochondrial dataset. Further, no model would be expected to fully recapitulate the evolutionary dynamics of the GPCR dataset, as these transmembrane proteins are subject to unique evolutionary constraints imposed by the membrane environment (Stevens and Arkin 2001; Spielman and Wilke 2013). We included the HIVb model as an example of a specialist model expected to fit poorly to the datasets used here, all of which differ from the data on which HIVb had been trained.

Although other analyses of protein evolutionary rates have opted to normalize rates by the gene-wide mean or median (Jack et al. 2016; Sydykova et al. 2018; Spielman and Kosakovsky Pond 2018) or convert rates to standard *Z*-scores (Pupko et al. 2002), we directly analyzed rates yielded by `LEISR` without any normalization. Because of this choice, we consider Spearman (rank) correlations when comparing MLEs. We adopt a rank-based test because MLEs are *relative* to the whole-protein rate, so the relative rank of MLEs is a more natural measure than the MLEs themselves. In addition, MLE distributions for a given gene are typically very skewed, with most rates falling below ~ 10 with a few outlying MLEs several orders of magnitude larger. We present many figures are on a log-log scale to capture the wide range of rate estimates.

### Gamma distributed rate variation has little effect on site-specific relative rates

We first compared, for each alignment, MLEs between each model’s inference, with or without +*G*. As expected, adding gamma variation resulted in increased tree lengths (Figures S1-2) compared to constant-rate models. However, the effect of +*G* on relative site rates was negligible. In Figure 1, we show how these inferences relate for single representative alignment from each of our four datasets. Rates track the *x* = *y* line of equality nearly perfectly across all datasets, with minor deviations generally appearing only at very high rates. Disagreements tend to emerge only for “fast” sites whose rates are difficult to estimate and have numerically unbounded confidence intervals (CI range ≥ 1000).

**Figure 1:**
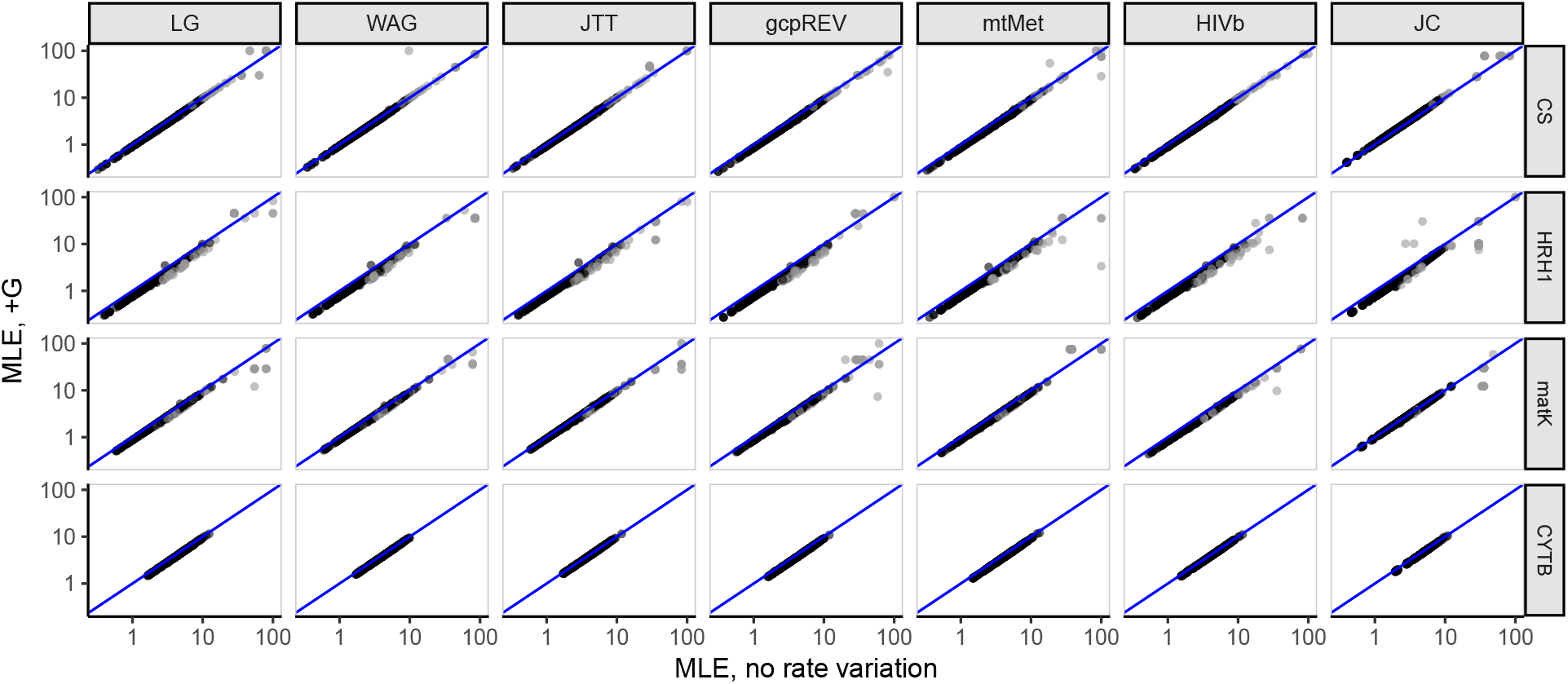
Relationship of site-specific MLEs inferred with a given model, with and without specifying +*G* in branch length optimization. Points in each log-log plot represent a single alignment site, and the line in each plot represents *x* = *y*. Representative alignments shown for enzyme, mitochondria, chloroplast, and GPCR datasets respectively are CS (citrase synthase), HRH1 (human histamine receptor 1), maturase K (matK), and cytochrome B (CYTB). Black points represent MLEs with reliable estimates, and grey points represent those with unbounded (range ≥ 1000) confidence intervals, for either axis. For visual clarity, all sites where MLE < 10^−8^, on either axis, have been removed from the figure.

This near-perfect agreement was consistent across all models considered, and all alignments examined here (Figure 2). The lowest measured correlation between model parameterizations was ρ = 0.924, and for 98% of comparisons, estimated correlation coefficients exceeded 0.99. We concluded that modeling rate variation during relative branch-length estimation has virtually no impact on the inferred site-specific rates MLEs. Consequently, we considered only rates inferred without +*G* for the remainder of our analyses.

**Figure 2:**
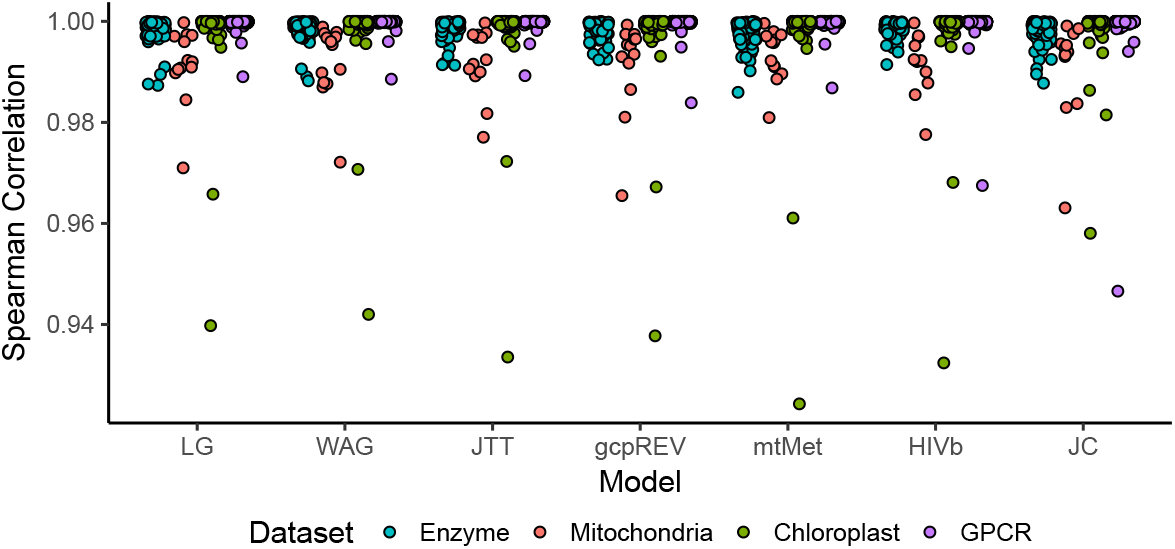
Spearman correlation coefficients of MLEs inferred with a given model, with and without specifying +*G* in branch length optimization. Each point represents the Spearman correlation between MLEs for a single alignment. Note the limited range for the *y*-axis.

### Models yield virtually identical relative rate inferences

We next assessed the extent to which the evolutionary affected relative rate MLEs at individual sites. In Figure 3, we show the relationship between LG MLEs and those inferred by all other models for four representative alignments across our datasets. A remarkably strong agreement was apparent for all model types (generalist, specialist, and equal-rates JC) and similarly for all datasets, regardless of taxonomy. Rate comparisons between LG and JC, however, did show somewhat more noise, although a clear rank correspondence for most sites was still present. Similar to the patterns observed in Figure 1, points which do not fall close to the *x* = *y* line in Figure 3 nearly always corresponded to sites with unreliable precise MLEs, i.e. sites with unbounded confidence intervals (CI range ≥ 1000).

**Figure 3:**
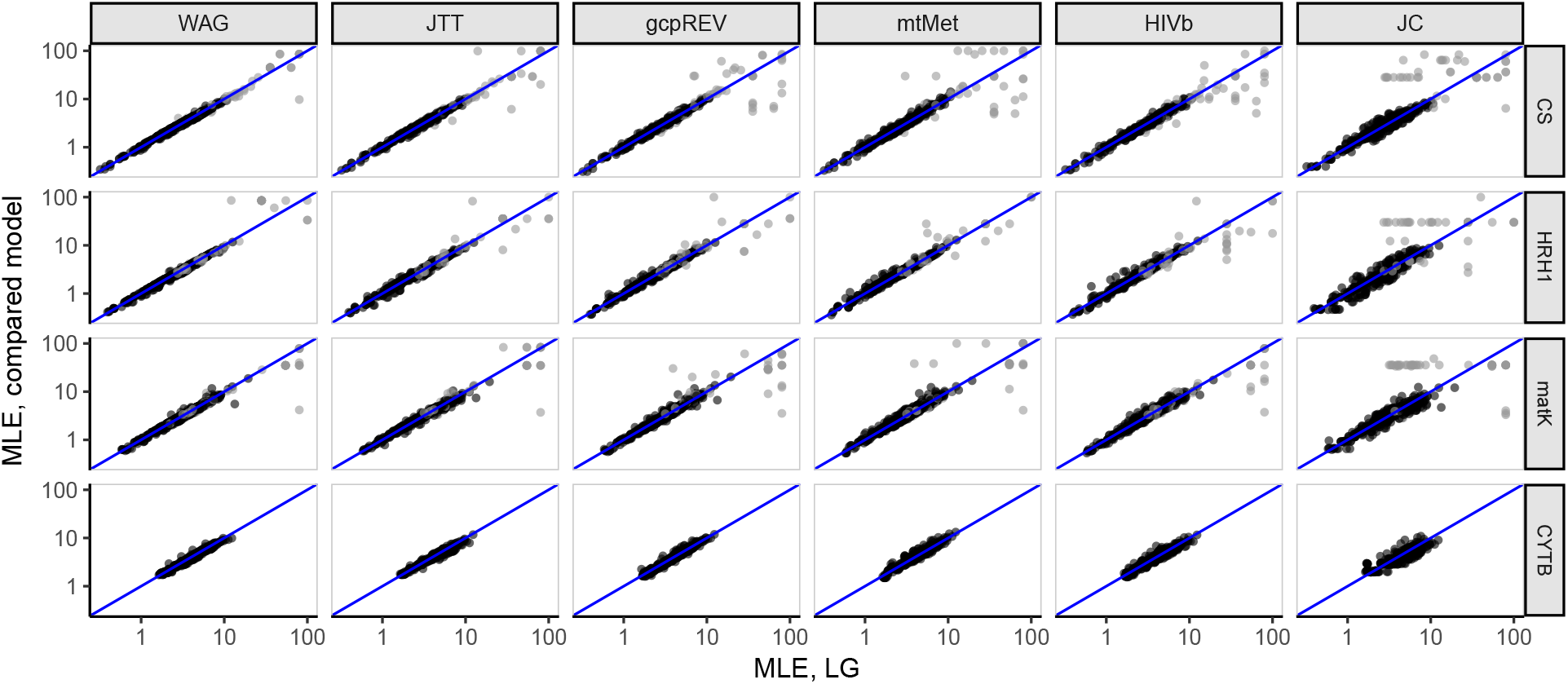
Relationship of inferred rates (“MLE”) between the LG and other models. Points in each log-log plot represent a single alignment site, and *x* = *y* line is also shown. Representative alignments shown for enzyme, mitochondria, chloroplast, and GPCR datasets respectively are CS (citrase synthase), HRH1 (human histamine receptor 1), maturase K (matK), and cytochrome B (CYTB). Black points represent MLEs with reliable estimates, and grey points represent those with unbounded (range ≥ 1000) confidence intervals, for either axis. For visual clarity, all sites where MLE < 10^−8^, on either axis, have been removed from the figure.

In Figure 4, we show Spearman correlations between all pairs of models, averaged across alignments for each dataset. Correlations were consistently high across all models and dataset types, generalist and specialist alike. For example, rates inferred on chloroplast with a chloroplast (gcpREV) model show nearly perfect correlations with rates inferred with a mitochondrial (mtMet) or HIV-specific (HIVb) model. JC was the only model with lower correlations with all other models, although the minimum averaged correlation was still high at *ρ* = 0.94 ± 0.024.

**Figure 4:**
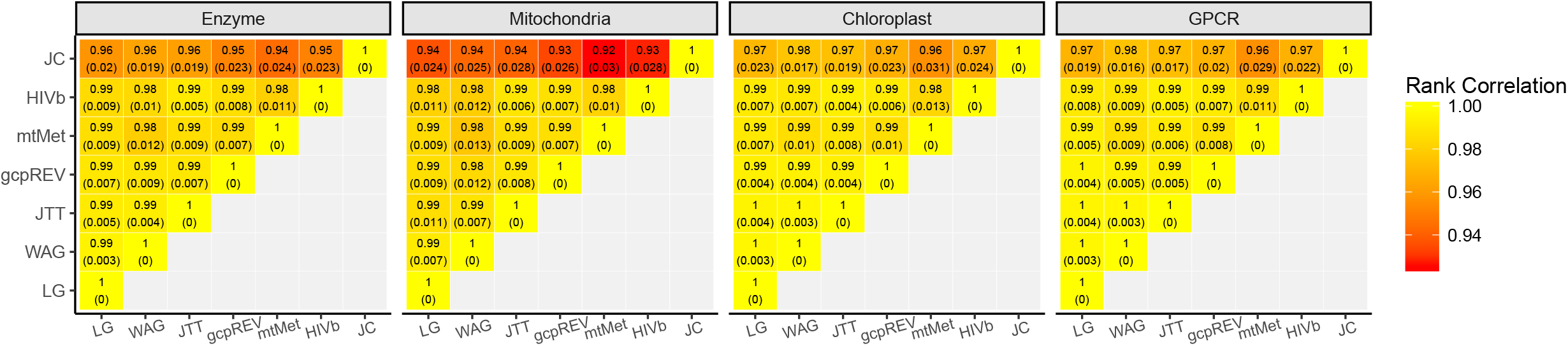
Heatmap of Spearman correlations between model inferences, considering only rates inferred without branch length heterogeneity. Values in each cell show the mean and standard deviation of Spearman correlations. Note that the heat scale ranges has a limited range between 0.95 and 1.

### Significant rate differences are infrequent but generally associated with JC

In spite of the near-perfect correlations among rates inferred with different models, relative rate estimates at individual sites are occasionally influenced by the choice of the model to a noticeable extent. For each individual site in all alignments (426,678 sites), we assessed the extent to which the approximate 95% confidence intervals (CI) from different models overlapped one another. For example, in Figure 5, we show five sites, each representing a different category of agreement or disagreement, from the enzyme citrase synthase alignment.

**Figure 5:**
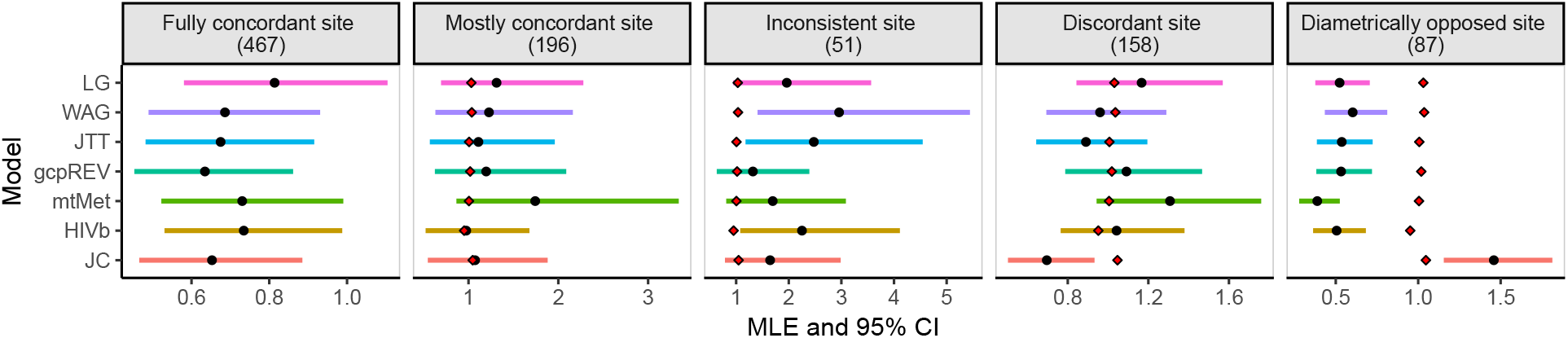
Relative rate inferences, bounded by 95% confidence intervals, for five sites from an enzyme alignment of citrase synthase sequences. In each panel, black points represent site MLEs, red diamonds (when present) represent the median gene-wide rate under the given model, and horizontal bars represent 95% CIs. See text for more details on site classifications.

1. A site was “fully concordant,” when the rate point estimate from any model fell within the 95% CI from every other model, indicating that the relative rate at this site was insensitive to model choice.
2. A site was “mostly concordant”, when the rate MLE from at least one model fell outside CIs from at least one other model (e.g., the MLE for mtMet fell outside the CI for HIVb), but all CIs included the median gene rate1 inferred from each model. In this case all MLEs did not significantly differ from the median rate, so a such a site would not be considered “interesting” for most downstream analyses.
3. An “inconsistent site” was a site where model inferences do not fully agree. At the example site in Figure 5, the MLE from the WAG model fell outside the CI for the gcpREV model; JTT and WAG models indicate that the site was evolving faster than the median site, while the other five models suggest that the site’s rate did not significantly differ from the median. However, all MLEs here displayed the same general trend of being larger than the median rate, and therefore sites like this would usually also not be considered “interesting.”
4. A site was “discordant” if at least one model (in the example case, JC), yielded a fully inconsistent relative rate estimate compared to most models. Specifically, the JC MLE for the example site’s rate was reliably below the median rate, whereas all other models yield MLEs that did not significantly differ from the median rate.
5. A site was “diametrically opposed” when, depending on the model, its rate was either reliably below or above the median rate.

The last two site classifications (discordant and diametrically opposed) reveal the most interesting sites for our purposes, in that different models inferred different and at times opposite levels of evolutionary constraint.

We queried all 426,678 sites across all alignments to determine how frequently the following scenarios occurred: i) fully or mostly concordant sites, ii) inconsistent sites, and iii) discordant or diametrically opposed sites. We found that only relatively few sites were impacted by the choice of model (Figure 6a), and the extent of discordance was influenced by tree length (Figure 6b). As tree length increased, the proportion of sites where models agree tended to decrease, while the proportion of sites where models disagree tended to increase. This observation was highly significant, as assessed with a linear model with proportion of sites as a response and a predictor as the interaction of tree length and model agreement represented by the three scenarios shown in Figure 6 (*P* < 2 × 10^−16^). In other words, the specific model chosen should have a larger affect on rate estimates for a more diverged alignment. By contrast, the specific model chosen may have virtually no impact on an alignment with relatively low divergence.

**Figure 6:**
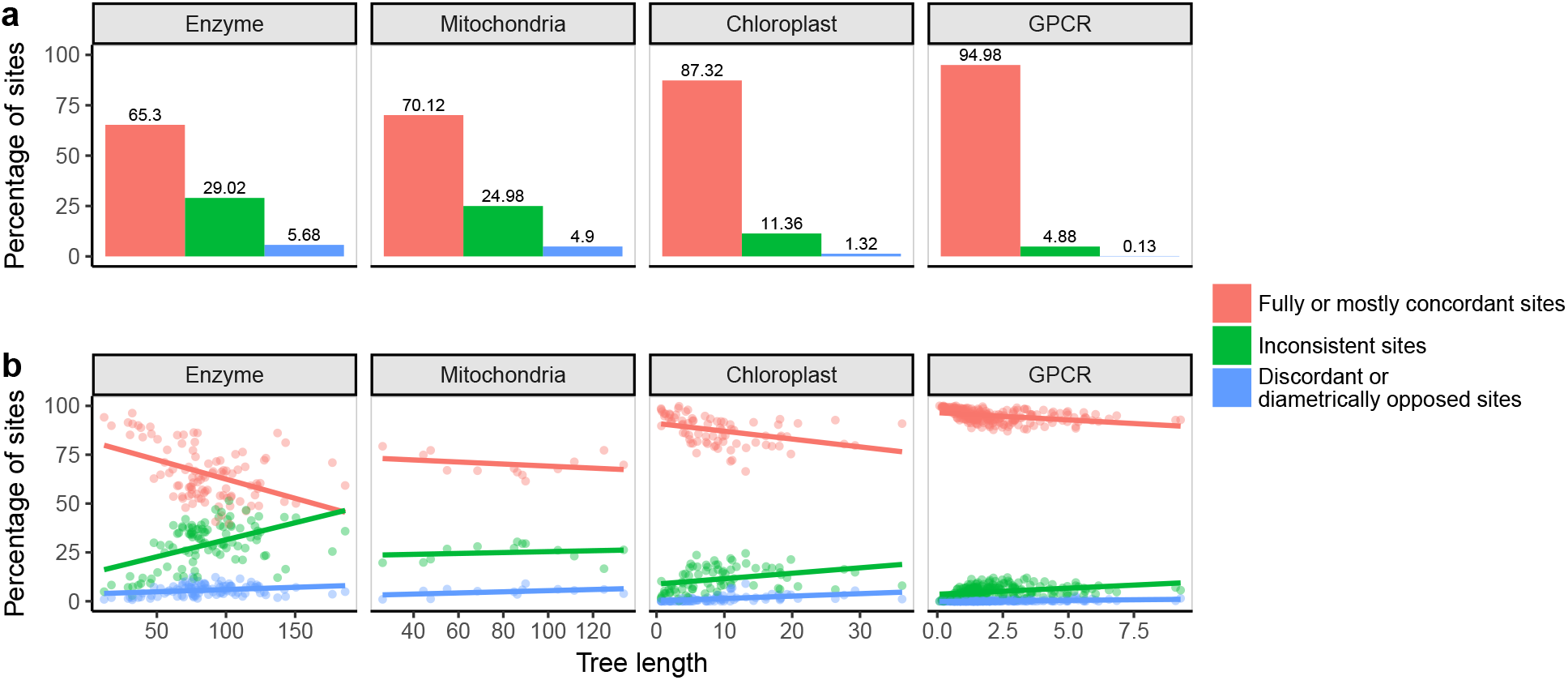
Classification of model agreement on individual sites. a) Average (over all alignments) percentage of sites, in each of our four datasets, which show different degrees of concordance among model inferences. b) Alignment-level model concordance as a function of tree length (substitutions / site / unit time), where each point represents a single alignment, and linear regressions are drawn as solid lines.

Only 118 sites (0.03% of all alignment columns in this study) belonged to the diametrically opposed category (Figure 5e), i.e., where at least one model reported a site’s rate as significantly lower than the median rate, while at least one other model reported it as significantly higher than the median rate. Strikingly, these 118 sites, found among 53 enzyme and 3 mitochondrial alignments, all followed the same pattern: JC was the outlying model, all JC MLEs were above the median rate, and MLEs from all other models were below the median rate.

We hypothesized that these sites represent fast-evolving residues where the fixed amino acids have relatively high exchangeabilities in empirical matrices. In such matrices, this fast-evolving site would appear to have a relatively low rate simply because the exchangeabilities would contain information that “should” be incorporated into the rate. By contrast, in JC, the exchangeabilities make no *a priori* assumption of fast evolution, and thus the rate parameter would be able to capture the truly high rate.

To probe this question, we directly counted the number of substitutions among each pair of amino acids at all sites, across all alignments. We employed HyPhy (Kosakovsky Pond et al. 2005) to count substitutions using joint maximum-likelihood (Pupko et al. 2000) under a specified amino-acid model (here, LG2) to reconstruct ancestral sequences. We tabulated substitutions under the principle of minimum evolution directly from the inferred ancestral sequences. Visual inspection of “diametrically opposed sites” revealed a high proportion of substitutions among the amino acids isoleucine (I), leucine (L), and valine (V), all of which have extremely similar biochemical properties and consistently high exchangeabilities across empirical protein models. We therefore determined the proportion of substitutions at each site, considering only sites which experienced at least one substitution, which were between either IL, IV, or LV ({*I, L, V*} substitutions)3. We found that diametrically opposed sites, and to some extent discordant sites, were strongly enriched for {*I, L, V*} substitutions (ANOVA *P* < 2 × 10^−16^, Figure 7A), with at least 56% of substitutions at the diametrically opposed sites occurring among I, L, and V. We additionally found that diametrically opposed sites tended to experience more substitutions (ANOVA *P* < 2 × 10^−16^, Figure 7B) relative to other site classifications. One interpretation for these results is that despite its poor fit to the data (based on information criteria or log likelihood), JC may uniquely capture sites of potential interest, where evolution rapidly occurs among substitutions with high model

**Figure 7:**
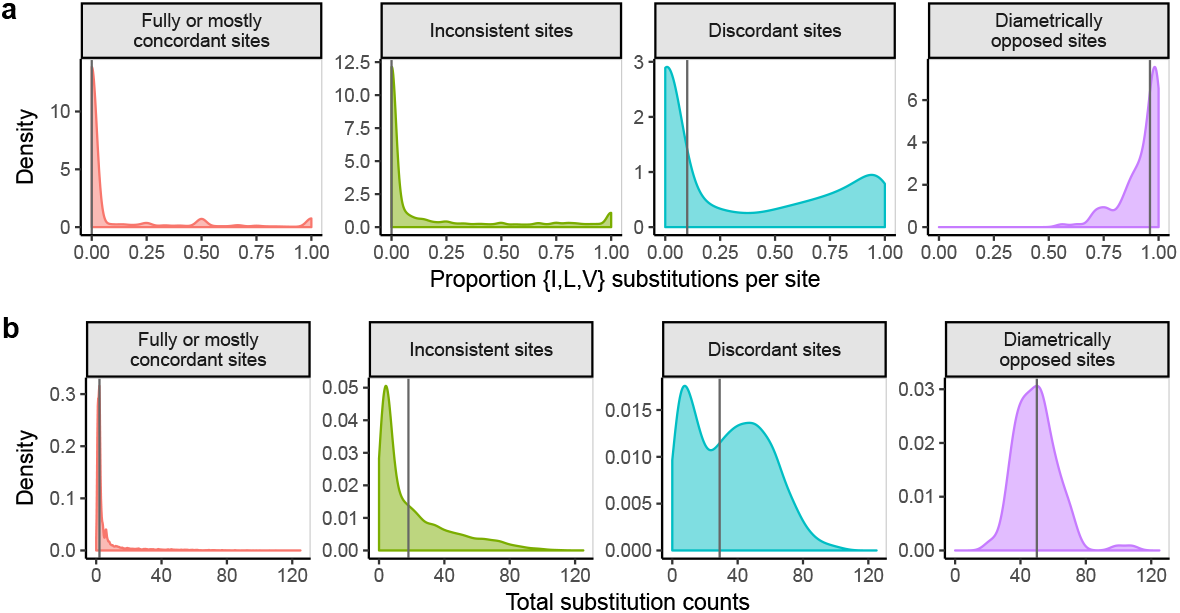
Distribution of substitution counts across sites. a) Proportion of {*I, L, V*} substitutions for all sites which have experienced at least one substitution, across site classifications as seen in Figures 5 and 6. The vertical line in each panel represents the median proportion of {*I, L, V*} substitutions. b) Total number of substitutions across site classifications, again only considering which had experienced at least one substitution. The vertical line in each panel represents the median total number of substitutions. Results in this figure were obtained using substitution counts under the LG model, but results are unaffected by the model chosen to count substitutions (see Figure S3 for analogous results obtained by counting substitutions under JC).

### Strong model fit differences are present in spite of high rate agreement

We determined which model would be preferred for each alignment using standard procedures in phylogenetic model selection (Posada and Buckley 2004). Specifically, we calculated the Akaike Information Criterion [*AIC* = 2(log *L* – *K*), where log *L* represents the log-likelihood and *K* represents the number of estimated model parameters)] for each model fitted to each alignment, and we ranked all models accordingly. We performed this model ranking separately for models with and without +*G*. Because any model under a given rate variation setting has the same number of parameters, other commonly-used information criterion measures such as small-sample *AIC* (*AIC_c_*) or Bayesian Information Criterion would yield the same results as *AIC* does here.

Considering only models without +*G*, the best-fitting model generally matched expectations given the scope of an alignment. All enzyme alignments were best fit by a generalist model (LG, WAG, or JTT), the majority of chloroplast alignments were best fit by the gcpREV model, and the majority of mitochondrial alignments were best fit by the mtMet model (Figure 8a, upper panel). GPCR alignments, which have no corresponding specialist model, were best fit by either a generalist model, mtMet, or HIVb (Figure 8a, lower panel). As expected, JC never emerged as a best-fitting model. These trends were broadly consistent with model selection results for +*G* models, with a few minor differences. Curiously, relatively more GPCR models showed a preference for HIVb than for a generalist model when +*G* was applied.

**Figure 8:**
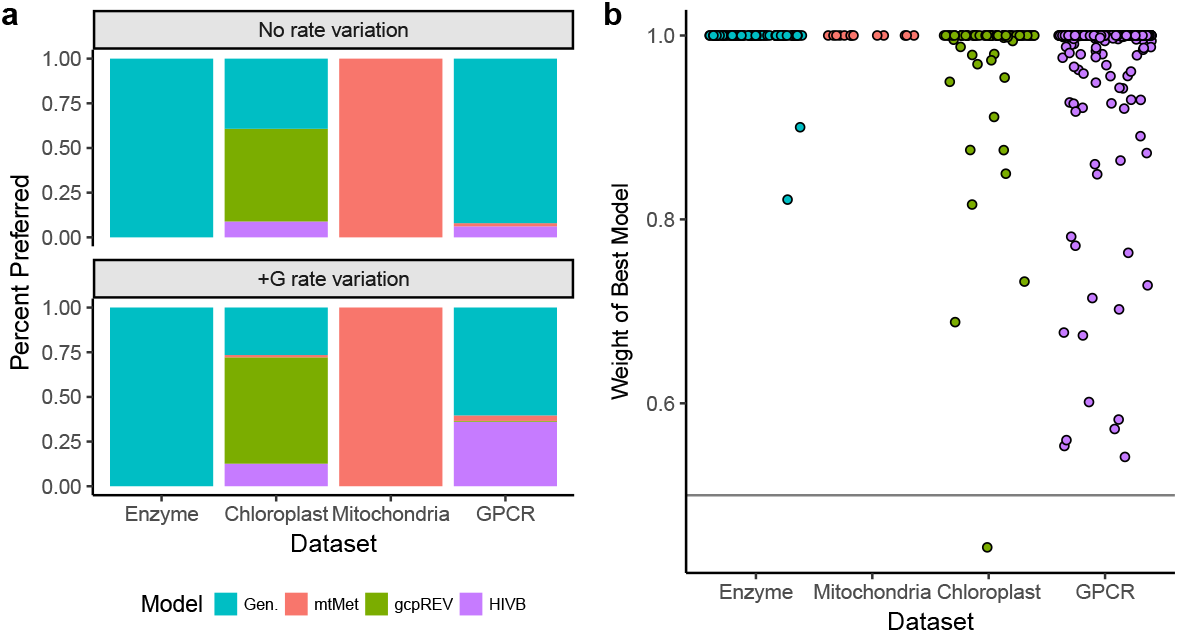
Model selection results. a) Distribution of preferred models, considering models without +*G* (top panel) and with +*G*. The legend abbreviation “Gen.” refers to one of the three generalist models (JTT, WAG, and LG) examined here. JC never emerged as a best-fitting model under either rate variation setting. b) Relative Akaike weight of top model for all alignments, grouped by dataset. The horizontal line is *y* = 0.5.

We determined the relative level of support for these preferred models (combining both +*G* models and models without rate variation), by calculated each model’s (relative) Akaike Weight. For each alignment, we calculated the weight *w* for each fitted model *i* as

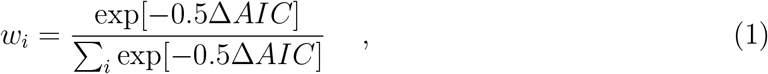

where Δ*AIC* = *AIC_i_* – *AIC*_min_, and *AIC*_min_ refers to the model with the smallest *AIC* value for the given alignment. For all but one chloroplast alignment, the best-fitting model (*AIC*_min_) was robustly supported with *w* ≥ 0.5 with the vast majority of weights at *w* ≈ 1 (Figure 8b).

We conclude that although virtually all alignments show strong model preferences measured by goodness-of-fit, empirically the models revealed broadly consistent and effectively interchangeable estimates of site-specific relative rates, with the possible exception of JC in certain circumstances.

## Discussion

We have investigated how the choice of empirical amino-acid evolutionary model affects inferred relative evolutionary rates in protein alignments. We conclude that for most sites in most alignments, the choice of substitution model has very little effect on the estimate. Even *a priori* poor choices, such as fitting a mitochondrial model to chloroplast data, failed to appreciably move the needle on site-specific relative rate estimates. More surprisingly, the model devoid of any biological realism (JC, Jukes and Cantor 1969), returned essentially the same estimates as the other models, with the exception of about one site in a thousand. While other applications of evolutionary rate have considered normalized and/or standardized rates (Pupko et al. 2002; Jack et al. 2016; Sydykova et al. 2018), which might emphasize agreement between models, our analysis found concordance between raw relative rates.

While rate inference appears to be quite robust to model choice, information theoretic criteria of model fit display very strong preferences towards a specific model of evolution (Figures 8), consistent with previous results and prior expectation. The goodness of fit improvement must therefore derive principally from features of the evolutionary process other than relative substitution rates at individual sites.

In general, the models examined here were developed for and have primarily been applied to questions of phylogenetic inference. Our results reveal both similar and distinct trends to those previously observed in the literature. For example, with respect to protein model performance, Keane et al. (2006) found that, in the context of phylogenetic tree inference, specialist models may not yield improved inferences relative to generalist models, even on specialist data. We have similarly shown that specialist and generalist models perform highly similarly when inferring relative evolutionary rates, for any given dataset. By contrast, Huelsenbeck et al. (2008) suggested that, again in the context of phylogenetic inference, no single protein model may be suitable for a given alignment. Our results instead suggest that, in the context of evolutionary rates, *any* protein model may be equally suitable for a given alignment, with the distinct possibility that they are all equally bad. However, we do emphasize that JC identified, albeit only very few, sites with salient signals of rapid evolution, where other more “realistic” models failed to identify these sites due to the high exchangabilities among substituting amino acids. As such, while JC consistently fits the data very poorly, it may in fact be a highly useful model that can identify rapid evolutionary toggling among highly-similar amino acids. Such evolutionary patterns can be highly biologically meaningful; for example, previous work has demonstrated that certain sites in HIV-1 experience strong selection pressure to undergo such amino-acid toggling (Delport et al. 2008). Therefore, it is possible that the simplistic equal-rates JC could be most useful for identifying certain types of selection pressures in proteins.

A careful analysis of properties of amino acid models including WAG, JTT, and LG by Goldstein and Pollock (2016) suggested that, due to the fact that these models strive to capture the propensities of an average protein, they effectively describe evolution neutrally, i.e. they don’t fit any particular protein especially well. Our findings on the application of such models to evolutionary rate inference echo the conclusion from Goldstein and Pollock (2016) that these models may not contain substantially different information about site-specific rates of protein evolution. However, our observation that rate inferences with JC, on occasion, uniquely deviated from models with unequal exchangeabilities implies that there is some difference between “averaged” matrices and neutral evolution, as JC would represent the exact neutral scenario of protein evolution, i.e., that any amino acid can be substituted for another with no selection preference or biochemical bias.

A wide variety of platforms have been popularized for selecting the best protein model in the context of both phylogenetic and evolutionary rate inference (Keane et al. 2006; Darriba et al. 2011; Ashkenazy et al. 2016; Lanfear et al. 2017; Kalyaanamoorthy et al. 2017). Model selection is considered a part of due diligence and good practice in these applications. However, an improvement in general goodness of fit may not translate into a quantifiable impact on the quantities of interest. For example, Spielman and Wilke (2015) found that, in the context of models of codon evolution (i.e. *dN/dS*-based models), *AIC* and *BIC* can positively mislead one to prefer a model with empirically worse rate estimates, while relatively poorly fit models in fact may produce the most accurate measures of selection strength.

Modeling assumptions commonly made for the sake of inferential tractability, e.g., site independence, using a fixed topology inferred from the same data for rate analysis, stationarity and time-homogeneity of the substitution process, are not biologically justifiable, but are tolerated because they produce biologically meaningful inferences. As George E.P. Box wrote, *“Since all models are wrong the scientist must be alert to what is importantly wrong”* (Box 1976). Our evidence is that choice of substitution model is “mostly harmless” (Adams 1979) for the purposes of site-level rate inference, and it is not necessary to chase the elusive best fit model for each protein alignment.

## Acknowledgments

This work was supported in part by grants R01 GM093939 (NIH/NIGMS) and U01 GM110749 (NIH/NIGMS).

## Footnotes

1 We considered the median gene-wide rate for this analysis, rather than the mean gene-wide rate, because difficult-to-estimate sites can yield inflated and highly outlying MLEs which overly bias the mean as a measure of location

2 We performed this procedure a second time with the JC model, results of which were indistinguishable from counts under the LG model (Figure S3)

3 Similar results were obtained for a more generic group of “highly exchangeable” residues (not shown)

